# Prediction of Algal Blooms in the Great Lakes through a Convolution Neural Network of Remote Sensing Data

**DOI:** 10.1101/450551

**Authors:** Karthik Srinivasan, Vikram Duvvur, Daniel Hess

## Abstract

Harmful algal blooms (HABs) are the proliferation of algae due to eutrophication and have severe repercussions to the ecological balance in many water bodies, due to the toxins the algae produce. Additionally, the identification and prediction of these HABs has been a challenge in the scientific community due to the interactions between both biological and physical processes that cause the HABs. Here, we used remote sensing data to bypass these issues; remote sensing data provides significant information about the coverage of chlorophyll which can be used to locate HABs. Using this indicator of HABs, we trained a Convolution Neural Network (CNN) to identify nine types of algal blooms, using 25 epochs of 900 images, which can predict algal bloom shapes with an 80 percent accuracy. This approach of HAB identification can easily be applied to other aquatic ecosystems where remote sensing data is present.

## Introduction

Algal blooms are a type of algal growth that can be dangerous to aquatic ecosystems in a variety of ways. Some blooms produce toxins capable of killing fish, mammals, and birds, and those toxins may also be harmful to humans (Landsberg, 2010). The death of the algae in the blooms can allow aerobic bacteria to consume all of the oxygen in their environment, leading to the formation of a dead zone. A dead zone is an area of water that has experienced hypoxia, or a loss of dissolved oxygen which has in turn led to the destruction of that area’s marine environment.

Algal blooms can also cause another problem, eutrophication. Eutrophication in a water body occurs when the level of nutrients, primarily phosphorus and nitrogen, increases significantly. The increased nutrient load causes a cascade of effects in which algal populations grow out of control and lead to a reduction in water quality and a destruction of the aqueous environment (Yang et al, 2008).

Table 1 shows a water quality classification table (Yang et al, 2008).

The criteria of surface water quality for lakes or reservoir (CNEPA, 2002)

| Items | Surface water quality classification |  |  |  |  |
| --- | --- | --- | --- | --- | --- |
|  | Class I | Class II | Class III | Class IV | Class V |
| Water temperature (°C) | Maximum week increase$\leq 1$; maximum week decrease$\leq 2$ | | | | |
| pH | 6~9 |  |  |  |  |
| DO (mg/L) | Saturation$\geq 90\%$ | $\geq 6$ | $\geq 5$ | $\geq 3$ | $\geq 2$ |
| COD<sub>Mn</sub> (mg/L) | $\leq 2$ | $\leq 4$ | $\leq 6$ | $\leq 10$ | $\leq 15$ |
| COD<sub>Cr</sub> (mg/L) | $\leq 15$ | $\leq 15$ | $\leq 20$ | $\leq 30$ | $\leq 40$ |
| BOD<sub>5</sub> (mg/L) | $\leq 3$ | $\leq 3$ | $\leq 4$ | $\leq 6$ | $\leq 10$ |
| TN (mg/L) | $\leq 0.2$ | $\leq 0.5$ | $\leq 1.0$ | $\leq 1.5$ | $\leq 2.0$ |
| NH<sub>3</sub>-N (mg/L) | $\leq 0.15$ | $\leq 0.5$ | $\leq 1.0$ | $\leq 1.5$ | $\leq 2.0$ |
| NO<sub>2</sub>-N (mg/L) | $\leq 0.06$ | $\leq 0.1$ | $\leq 0.15$ | $\leq 1.0$ | $\leq 1.0$ |
| TP (mg/L) | $\leq 0.01$ | $\leq 0.025$ | $\leq 0.05$ | $\leq 0.1$ | $\leq 0.2$ |
| Chlorophyll a | $\leq 0.001$ | $\leq 0.004$ | $\leq 0.01$ | $\leq 0.03$ | $\leq 0.065$ |
| Transparency (m) | $\geq 15$ | $\geq 4$ | $\geq 2.5$ | $\geq 1.5$ | $\geq 0.5$ |
| *Escherichia coli* (L<sup>-1</sup>) | $\leq 200$ | $\leq 2000$ | $\leq 10000$ | $\leq 20000$ | $\leq 40000$ |

DO: dissolved-oxygen; COD<sub>Mn</sub>: Chemical oxygen demand by K<sub>2</sub>MnO<sub>4</sub> oxidation method;
COD<sub>Cr</sub>: Chemical oxygen demand by chromium oxidation method; BOD<sub>5</sub>: Biological oxygen demand; TN: Total nitrogen; TP: Total phosphorus

Table 2 shows a comparison of different types of aquatic ecosystems and their water quality (Yang et al, 2008).

**Table 2.**
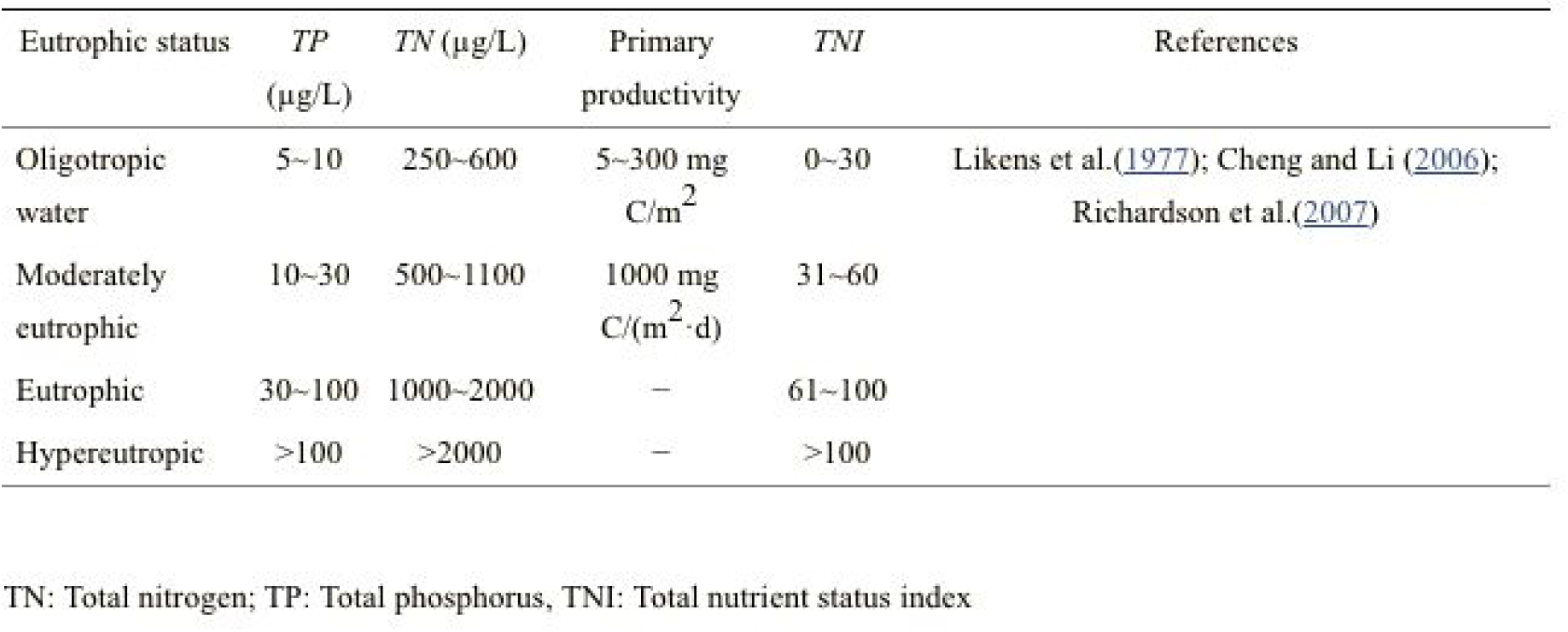
The burden value of N and P in various eutrophicated water

| Eutrophic status | TP<br>($\mu\text{g/L}$) | TN ($\mu\text{g/L}$) | Primary<br>productivity | TNI | References |
| --- | --- | --- | --- | --- | --- |
| Oligotrophic<br>water | 5~10 | 250~600 | 5~300 mg<br>$\text{C/m}^2$ | 0~30 | Likens et al.(1977); Cheng and Li (2006);<br>Richardson et al.(2007) |
| Moderately<br>eutrophic | 10~30 | 500~1100 | 1000 mg<br>$\text{C}/(\text{m}^2 \cdot \text{d})$ | 31~60 | |
| Eutrophic | 30~100 | 1000~2000 | — | 61~100 |  |
| Hypereutrophic | >100 | >2000 | — | >100 |  |

TN: Total nitrogen; TP: Total phosphorus, TNI: Total nutrient status index

A variety of conditions may cause HABs. The main factors are temperature, turbidity, nutrient levels, wind, water currents, and light levels. Of these factors, nutrients - most notably phosphorus and nitrogen - are the single most significant cause of HABs. Nutrients, primarily nitrogen and phosphorous, that because of their quantity limit algae growth, are called limiting nutrients. When the levels of the limiting nutrients in the water build up past acceptable levels algal blooms are often caused. Runoff from fertilizers used on lawns and in agriculture is one of the main ways that nutrient levels increase to the point where they are causing algal blooms.. When it rains, these fertilizers are washed into lakes, seas, rivers, and oceans, where they wreak havoc. The increase in nutrient levels, especially limiting nutrients, allow the algal population to significantly proliferate.

The same factors that precipitate algal blooms may be used to predict their occurrence, shape, and severity, if those blooms occur between May and September. But by using a different set of tools - specifically, supervised machine learning and satellite imagery - it is possible to predict the likely occurrence and distribution of algal blooms a month in advance over large areas. Supervised machine learning is the use of an artificial intelligence that creates an algorithm based on classified, or organized and labeled, data to, in this case, predict algal blooms. The software learns from the example, and builds its own algorithm based solely on the inputs and outputs. The satellite imagery showed the micrograms per liter of chlorophyll in the Great Lakes.

Chlorophyll monitoring is essential for the prediction of algal. Satellite remote sensing is the best method for identifying HABs in a large area. Remote sensing is more advantageous than any other form of HAB identification (Wei et al). Because remote sensing of chlorophyll enables the large scale detection of HABs we utilized it for our predictor.

If researchers could use satellite images to measure nitrate and phosphate levels, algal blooms could be forecasted earlier and with greater precision than they are today. But using current sensing technology, these ions are not visible from space. The amount of chlorophyll in water, however, can be used as a proxy. Photosynthetic activity is easily detected by satellite imaging. We chose to use chlorophyll levels for two reasons: they are easy to measure from space, and they provide a good, early indication of potentially problematic algal blooms. Research has shown a strong positive empirical relationship between the number of bacteria and the amount of chlorophyll in both freshwater and marine environments. The difference between freshwater and marine environments is negligible as the linear regression equations are statistically indistinguishable (Bird and Kalff).

Because of chlorophylls and bacteria relationship, high levels of chlorophyll correlate to HABs. We therefore can use the chlorophyll levels to predict the HABs. Our accuracy for identifying HABs is 80 percent.

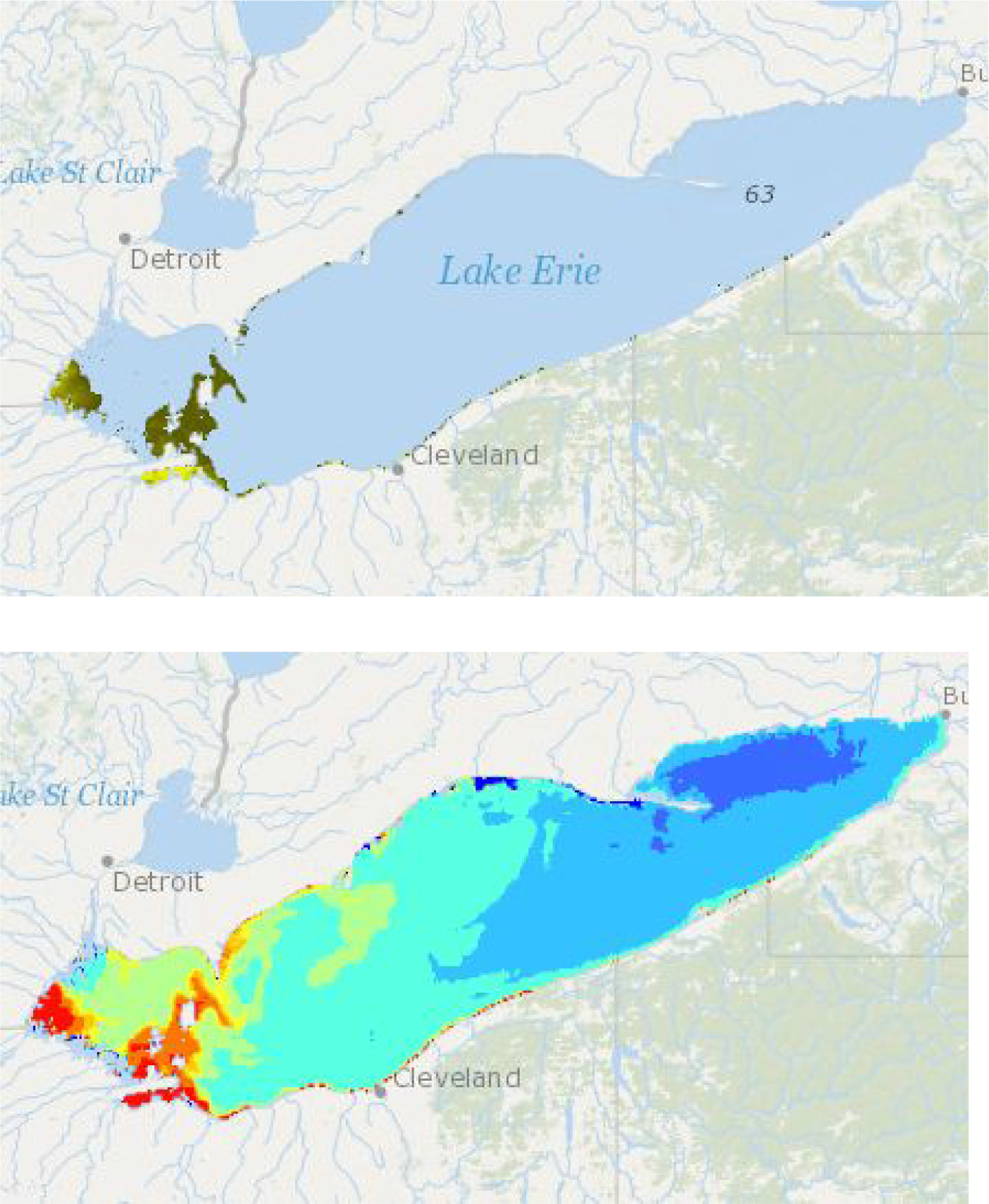
A comparison of chlorophyll and HABs in Lake Erie on August 23rd 2018.

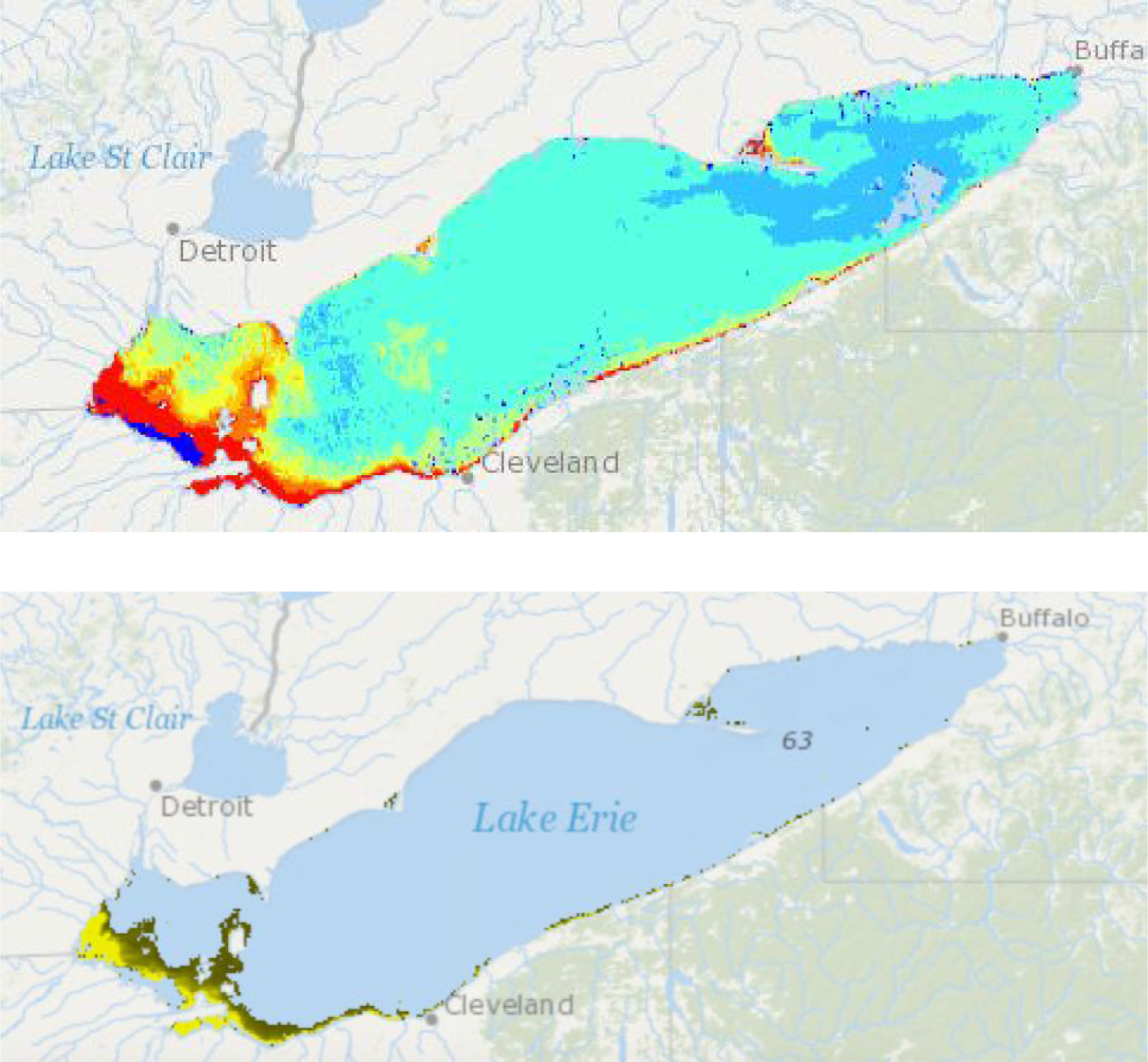
A comparison of Chlorophyll and HABs in Lake Erie on July 4th 2018.

## Methods

Machine learning is a type of artificial intelligence (AI) in which a training set is used to teach the AI without the need for it to be explicitly programmed. We needed a neural network capable of identifying several types of common algal bloom shapes in the form of images, so we decided to create a Convolution Neural Network. A neural network is made up of multiple “layers”, each of which serves a different purpose and is composed of nodes that contain the logic of the network. We chose to make our neural network a back-propagation network, because the network can evaluate the error and propagate it back through earlier layers to get more accurate classifications of algal bloom shapes. The results are evaluated by loss functions, which reward higher accuracies with lower loss values. Ultimately we want to place an image into one of 11 categories: nine shapes of algal blooms, plus water and land. These categories are known as classes, hence the name multiclass identifier. We coded the project in Python 3.4 using the Keras and TensorFlow libraries, but the methods will be described in pseudocode.

### Neural Network Structure

We decided to use a specific type of machine learning model called a Convolution Neural Network because it is frequently used for multiclass classifiers for visual images. To prevent overfitting of the data and reuse of data, we started by setting up a convolution layer that applies 32, 3×3 filters to each image. To reduce the difficulty of classifying images, we simplify the upcoming layers by setting up a 2×2 pooling layer. This pooling layer aggregates information from the previous layer. After the pooling layer, we flatten the data into a one-dimensional single vector to ensure the next layer analyzes all the data from the previous layers. We then create a hidden layer with 128 leaky rectifier linear unit nodes, because this fully connected layer is how we can find the non-linear combinations of features extrapolated from the previous layers. We determined 128 hidden nodes would be an ideal amount of nodes, because we made several neural networks and used 200 training and 100 validation images for testing powers of two hidden nodes from 2^2^ **to** 2^10^, **and** 2^7^ **had the highest accuracy**. Leaky rectifier nodes ensure that the value of the nodes do not get stuck at zero and cripple the neural network’s signature back propagation. The final output layer is an 11-unit sigmoid node layer, which gives 11 values ranging from zero to one depending on how closely it matches it matches each class, with “1” being an exact match and “0” being not close at all. The 11 classes are nine types of algal blooms, one of land, and one of water. The purpose of the sigmoid function allows the 11 classes to be easily compared, so that the node with the highest value corresponds to what the image most likely contains. Then we compiled the network with the Adam loss function, which is used to evaluate how effective the weights of the current iteration of the model are. We chose the the Adam optimizer because it is considered to be computationally efficient (Kingma and Lei Ba, 2015).

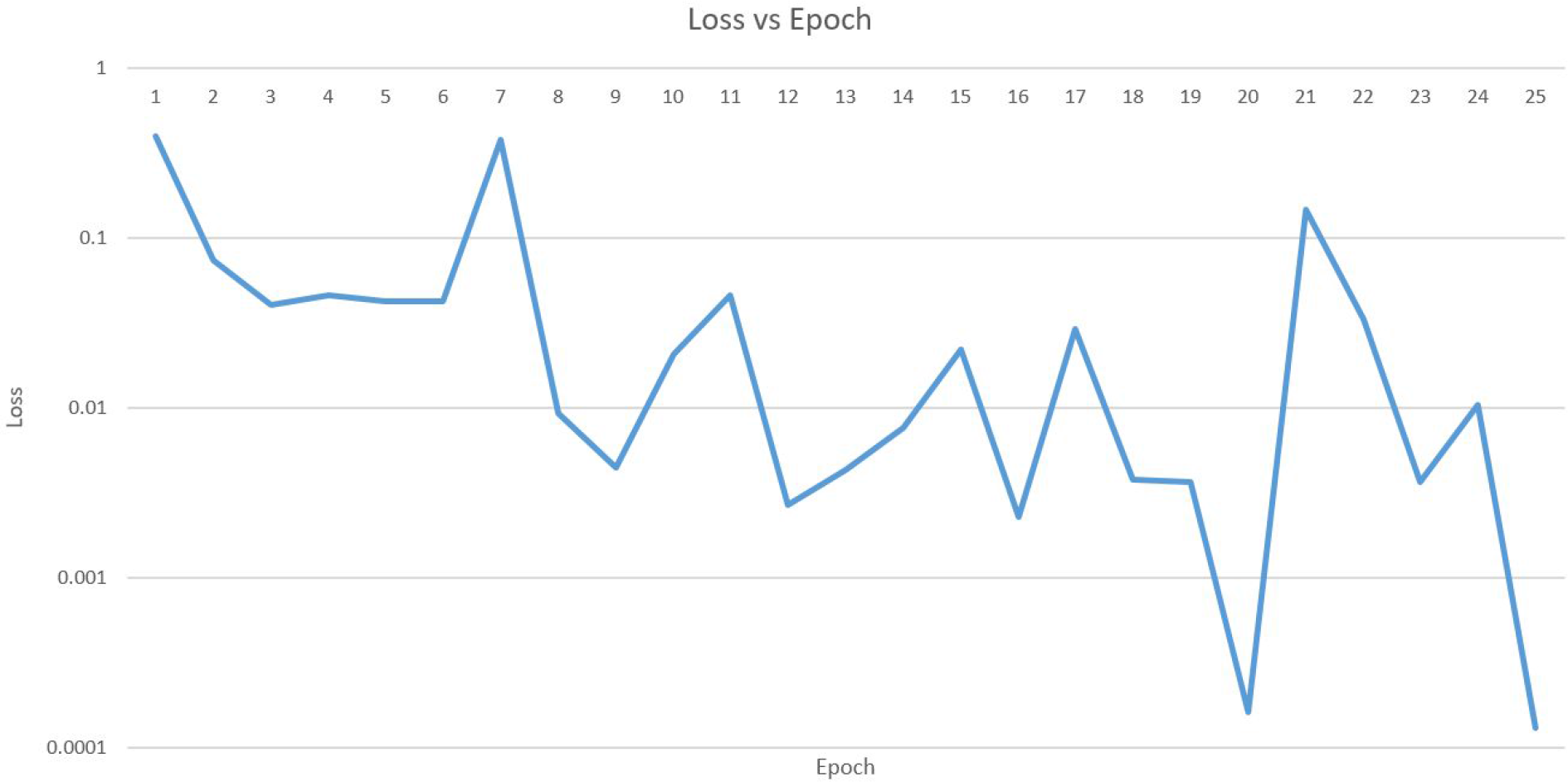
Pictured above is the graph of the loss of our model versus its epoch through the data on a logarithmic scale.

### Training Images and Labeling

We used 650 images from the Great Lakes Remote Sensing Chlorophyll-A map, created by the Michigan Tech Research Institute, to select the training examples for our Convolutional Neural Network. The images were taken from 2010-2015. We used 50 to 60 images for each of the 11 categories. We decided on the number of epochs by running a grid search ranging from 1 epoch to 40 epochs by multiples of 5, and the 25 epoch neural network had the highest accuracy for our 100 picture validation set. Our convolution layer allowed us to go over the same 650 images multiple times, effectively increasing our total training set to over 16,000 images.

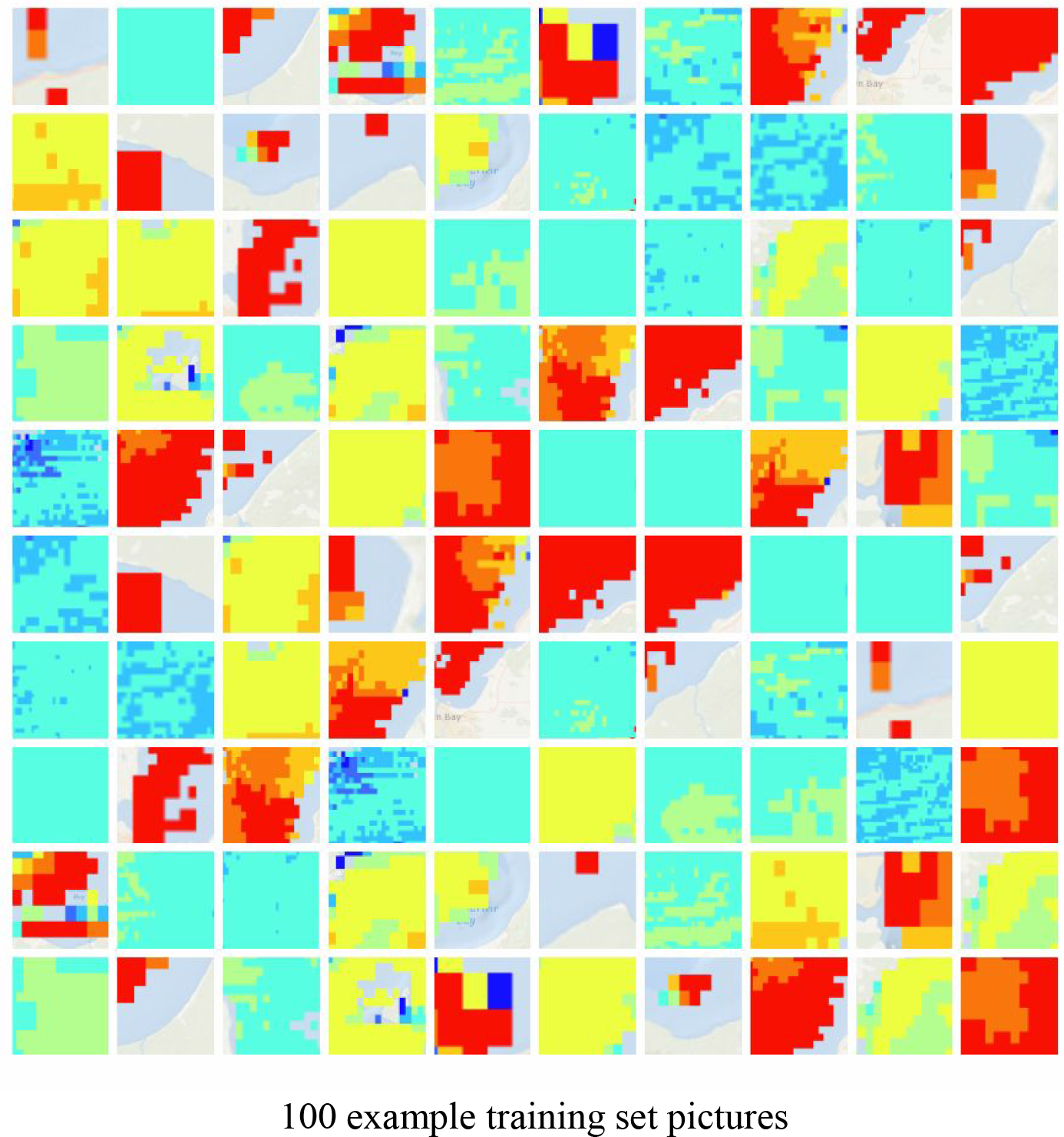
100 example training set pictures

### Prediction

Given a Chlorophyll-A map, the program first cuts it up into 32 by 32 pixel squares. It then runs the identification algorithm on all of the 32 by 32 squares. It then predicts the next stage of the algal bloom according to the flowchart below, and replaces the square if it has a progression on the chart. Finally, the image is stitched back together into one output image.

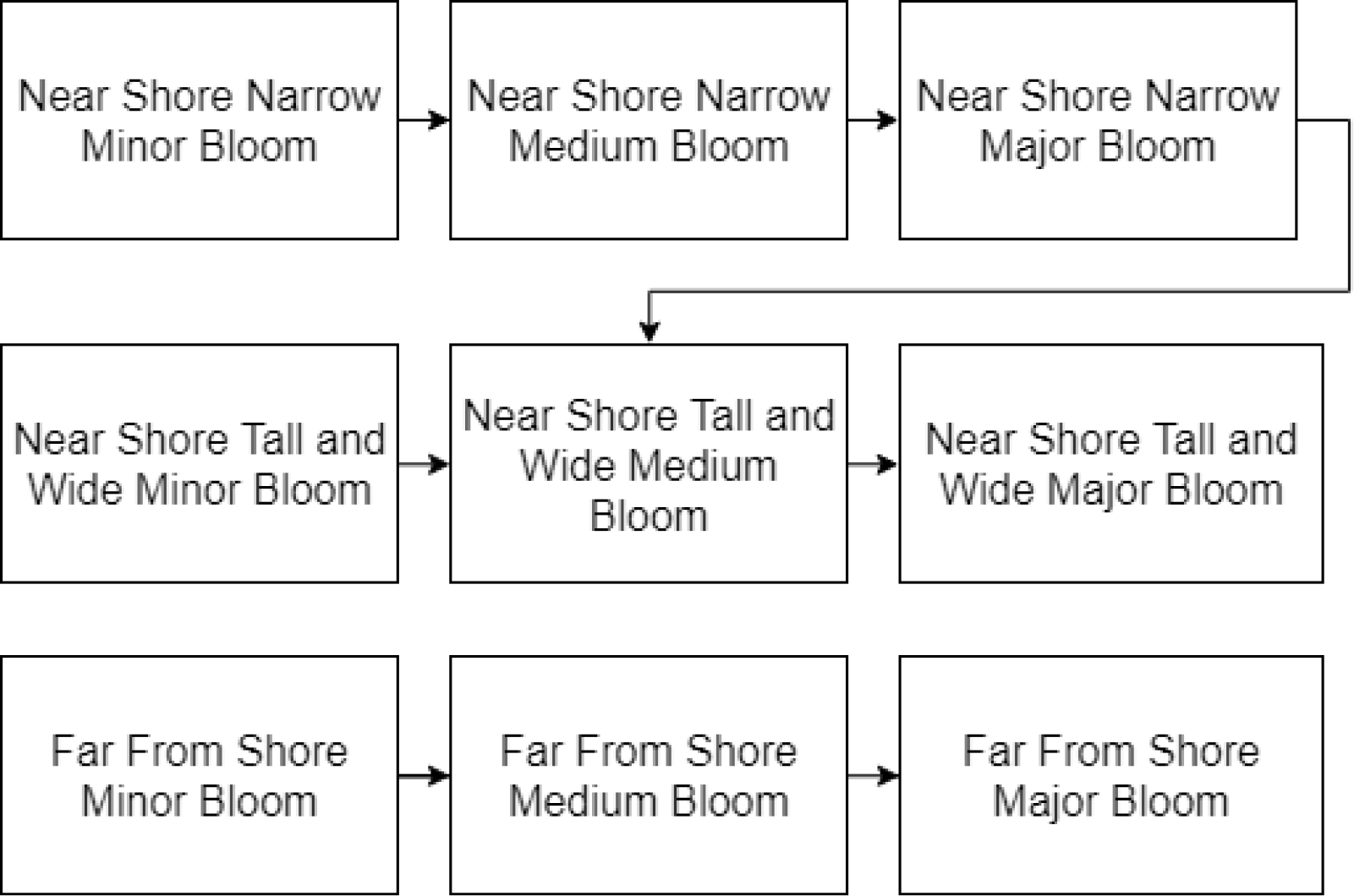
This is a flow chart of how the prediction algorithm predicts the algal bloom growth based on the previous bloom.

## Discussion

Our algal bloom predictor identification accuracy was 80 percent across the nine types of algal blooms, compared to 9.1 percent identification rate from the random assignment of classes. The model will be able to provide an image predicting where and how algal blooms are likely to grow in the future. The predictive power of our software can be best leveraged to mitigate the the damages to local businesses and wildlife from harmful algal blooms. That is because algal control methods taken in advance of a bloom are the most effective way of preventing and minimizing their effects.

Algal blooms have become a greater threat to natural environments in recent years, and predicting and monitoring them is essential to mitigating the damage they may cause. This approach was advocated in 2001 in a report to Congress by the National Oceanic and Atmospheric Administration and the National Sea Grant College Program that was created by the Woods Hole Oceanographic Institution. The monitoring method would need to allow local, state, and federal groups to work together. It would have to have early warning capabilities and accurate forecasting. It would have to predict bloom occurrence, development, and location. This type of prediction would enable the deployment of realistic mitigation strategies, such as reducing nutrient runoff from yards or farms. By doing this, human health and economic impacts could be reduced. Our software provides the exact kind of service that was called for in 2001. It is real time, predictive, accurate, and would enable both preventative and mitigatory steps to be taken to combat algal blooms.

Selected samples of water eurtriphication occurrences in lake, reservoir, estuary and river in the word

| Water body | Nutrient (N, P) concentrations | Phytoplankton biomass | History of eutrophication | References |
| --- | --- | --- | --- | --- |
| Erie Lake, USA | TP: 115 µg/L | Chlorophyll a: 58 µg/L | Blu-green algae bloom between 1965 and 1979; Declined quality from 1995 | Reutter, 1989; Maggie, 2004 |
| Okeechobee Lake, Florida, USA | TP: 50–100 µg/L | N and P are limitors for phytoplankton | Eutrophic at present | Schelske, 1989 |
| City Park Lake, Louisiana, USA | TN: 682 µg/L, TP: 330 µg/L | Chlorophyll a: 35.1 µg/L | Eutrophic (2000–2001) | Ruley and Rusch, 2002 |
| Lugano Lake | TP: 140 µg/L | Phytoplankton dry weight: 7–16 g/m<sup>2</sup> | Strongly eutrophic from 1960s | Barbieri and Simona, 2001 |
| Danish lakes | TN: 2.4 mg/L, TP: 0.37 mg/L | Chlorophyll a: 73 µg/L | Highly eutrophic | Jeppesen et al., 1999 |
| Pamvotis Lake, Northwest Greece | TP: 0.11 µg/L, NH<sub>4</sub><sup>+</sup>: 0.25 mg N/L, NO<sub>3</sub><sup>-</sup>: 0.56 mg N/L | – | Cultural eutrophication over the past 40 years; Eutrophication at present | Romero et al., 2002 |
| Berg River, South Africa | TN: 217 mg/L, TP: 70 mg/L | – | Hypertrophic | de Villiers, 2007 |
| Chivero Lake, Zimbabwe | TN: 0.3–8.4 mg/L, TP: 1.01–5.01 mg/L | Chlorophyll a: 18.02–22.48 µg/L | Highly eutrophic since 1980s | Nhapi, 2004; Ndebele and Magadza, 2006 |
| Taihu Lake, China | TN: 2.56–4.5 mg/L, TP: 0.25–0.35 g/L | Algae biomass: 2.7–6.4 mg/L | Class I/II in the early 1960s; Class IV in the mid-1990s; Now, inferior Class V | Ye et al., 2007; Jin et al., 2006 |
| Songhua Lake, China | TN: 1.14–1.98 mg/L, TP: 0.038–0.102 mg/L | Algae density: 210.84×10<sup>4</sup>–432.68×10<sup>4</sup> cell/L |  | Wang et al., 2004 |
| Dianchi Lake, China | TN: 2.13–8.27 mg/L, TP: 0.33–0.59 mg/L | – | Class III in the early 1970s; Now, inferior Class V | Guo and Sun, 2002 |

**Table 3** shows selected samples of water eutrophication occurrences in selected lakes, reservoirs, estuaries and rivers around the world (Yang et al, 2008).

Algal blooms are based on the water conditions that are most often tracked in real time through buoys in the water or with water sample testing. (Yang et al, 2008). This is very constraining on predicting the size, scope and nature of algal blooms, because such an approach requires that scientists have direct access to a specific area of a body of water. The buoy method also limits the area for which an accurate forecast can be made because researchers must have access to that particular area of water or be able to place sensors in those areas. In addition, it greatly increases cost because scientists have to physically collect water samples or place sensors. By utilizing existing data that is already being collected and employing machine learning, we sidestep the normal costs that are associated with forecasting algal blooms. We do not need to be at the site of the algal bloom, and we do not need to physically collect water samples, which are costly factors. All the work done with our prediction system happens on a computer and can be done by anyone.

According to the Environmental Protection Agency, methods for combatting algal blooms are grouped into three main categories: mechanical; physical or chemical; and biological. Mechanical control methods involve pumps, skimming, and barriers. Aerator pumps de-stratisfies water by pumping air through the water, however each device has limited range. Another type of pump, known as a mechanical circulator, pumps water from the surface down to the bottom to disrupt the stratification of water. This suffers from the same drawback as aerator pumps, and some forms of algae prefer a turbulent environment. Algal Blooms often form visible blooms on the surface, which can be removed through the use of surface skimmers similar to those used in oil spills. The major disadvantage of this technique is that it requires the bloom to be in a late developmental stage, which mean most of the environmental devastation has already been caused. Another type of physical control method is use of ultrasound. An ultrasonic device can emit certain frequencies of sound waves that destroy cyanobacteria cells, however it can also harm non-harmful green algae. More research is required to determine the true effectiveness of this method. Physical or chemical control relies on the use of chemicals or mineral compounds to combat algal blooms. Algaecides such as potassium permanganate, chlorine, and lime have been historically used to combat blooms, and thus their effects are thought to be well known, but certain algaecides can kill aquatic wildlife. Coagulants and flocculants sink cyanobacteria cells and nutrients to the bottom of the water column respectively, but both are only as effective as the depth of the water (US EPA). Biological controls are often the most complicated. They rely on the use of organisms or pathogens, such as viruses, bacteria, parasites, zooplankton, and shellfish, that are able to kill or lyse HAB cells (Wei et al).

HABs are dangerous not only to the aquatic systems in which they occur, but also to the humans who use those systems. Because algae are eaten in large amounts by many aquatic animals and help to form the basis of the trophic pyramid, dangerous algae can cause a build up of toxins in the fish and other aquatic organisms that humans consume. HABs have a significant impact on the aquatic resources and can affect how resources are used. The best example of the human impact of harmful algal bloom is the death of humans caused by the consumption of shellfish. Shellfish consume the toxic HABs, and the toxins build up inside of the shellfish. When humans consume these tainted shellfish, they can then become sick and die (Wei et al). The direct human impact of HABs make it essential that our software is utilized to reduce and combat algal blooms.

Algal blooms do not just present a health risk for humans; fish die-offs are a common side effect of algal blooms. When algae flourish in an aquatic environment, they rapidly use up all the dissolved oxygen in whatever area they are growing in. Once they consume oxygen, very little oxygen remains available for other organisms, and mass die-offs may follow. In addition, algal blooms almost always occur during times of warmer temperatures. When water is warmer, it holds less dissolved oxygen naturally, meaning that there are often two factors depleting levels of dissolved oxygen. Due to the shortages of oxygen caused by these two factors, fish can often suffer die-offs (Yang et al, 2008).

On top of the health risks caused by algal blooms, they also may impose significant economic costs. HABs have a massive economic impact, both directly and indirectly, in the areas where they occur. An outbreak of paralytic bacteria in the Northeast, That was caused by HABs, had an estimated cost of six million dollars (Shumway 1988). In North Carolina between 1987 and 1988, a four- to six-month period of red tide had an estimated cost on the community of 25 million dollars (Tester and Fowler 1990). A 1991 domoic acid and amnesic shellfish poisoning outbreak in Washington State had a huge impact on the entire community. It affected tourism, fisheries (mostly oyster), resulting in losses valued at between 15 and 20 million dollars. An outbreak in North Carolina of Pfiesteria piscicida and Pfiesteria between 1995 and 1996 caused massive damage. Millions of fish, including the commercially fished menhaden, died due to toxins and or secondary infections (Burkholder and Glasgow 1997). Another outbreak of Pfiesteria or Pfiesteria-like organisms caused an estimated 43 million dollars in damages to the Chesapeake Bay region. It resulted in a public outcry and several reported cases of illness (Sieling and Lipton 1998). The economic impact of HABs is substantial and affects many parts of the world. Because of this, it is essential that we create new prediction methods like our algorithm so that control methods can be better utilized (Bushaw-Newton, K.L. and Sellner, K.G).

Though marine coasts experience numerous algal blooms, the issue also is significant in the Great Lakes. Federal agencies have begun to take note of the problem in the Great Lakes, especially in Lake Erie, where the blooms are particularly common. Recent research has shown a link between health problems in residents of the Great Lakes area and blooms of blue-green algae, otherwise known as cyanobacteria. Blue-green algae blooms occur frequently in the U.S. They are often associated with eutrophication and the nutrient enrichment caused by runoff that causes the blooms. Blue-green algal blooms usually form on the surface of water and occur most commonly during the warmest times of the year. They are frequently a source of annoyance for people who participate in boating, fishing, and swimming. They also often cause flavor and odor problems in water and at water treatment plants (Wei et al).

Our predictor is meant to work alongside the methods of algal bloom detection and prediction. By creating a method of algal bloom prediction that works off of satellite imagery we enable more exact methods of detection and data collection to be more accurately placed where they can be most effectively used. Our predictor provides a preview of what algal blooms might look like so that measures can be taken to verify and then combat the algal blooms.

## Results

### Prediction of only HABs

The cyclical nature of the HABs throughout the seasons of the year hints to the fact that HABs are, in fact, predictable. The correlation between a predictive algorithm for HABs and the HABs that occur in the future is quite pronounced (Muttil et al, 2005). The satellite imaging data captured the amount of photosynthesis occurring. We were able to use this data to predict HABs, as opposed to merely photosynthetic activity, due to the correlation aforementioned. The reasoning behind this logic is sound. Although the images are marked as measuring all photosynthetic activity, the power of the satellite images would not be nearly enough to reach the activity of photosynthetic plants in the benthic region of deep, oligotrophic lakes such as those in the Great Lakes. As a result, the photosynthetic activity being depicted in the images must be in the epilimnion. Apart from those that are able to grow in the littoral zone, the only photosynthetic activity that would be occurring in the epilimnion would be that of algae, and any algal colony large enough to be significant in the satellite images suggests that a bloom is occurring.

### Other Methods

HABs are a major problem all over the world. In the Great Lakes, Lake Erie, in particular, has suffered from blooms. Our prediction method enables prediction of algal blooms not just in the Great Lakes, but in all bodies of water. A widespread literature review was performed to understand the HAB problem in Lake Erie. The current methods to forecast HABs all over the world and, specifically, in Lake Erie were examined. An extensive literature review and analysis was performed on the possible variables for forecasting HABs. Two forecasting methods - CART, classification and regression tree, and ANN, artificial neural network - as well as two training periods and two input variable selection methods, nutrient loading period and Spearman’s rank correlation were used. The Spearman rank correlation describes how well two variables’ relationship can be explained with a monotonic function. A monotonic function is a function that is always either entirely non-decreasing or entirely non-increasing. For the nutrient loading period selection method, only one set of input variables is used for forecasting. The Spearman selection method, on the other hand, examines more variables than the nutrient loading period, considering up to 28 different averaging periods and lag times for each considered variable. First, the CART forecasting models were tested with both classification methods, a 3-class and 5- class system resulting in the 3-class system being selected. The CART models were then created for both methods and training periods.

Initially, when using the first training period of 2002 to 2011, the loading period method showed better precision in forecasting HABs when compared to the Spearman selection method. When the training period was increased to 2013, both methods showed an improvement in the overall accuracy, with Spearman having an 8.9 percent improvement, and loading period a 5.4 percent improvement. However, with the extended training period, the loading period decision trees for August and September showed a slight increase in precision over the Spearman method and the Spearman method being slightly more precise in October. After the CART models, the ANN models were created and analyzed for both selection methods and training periods. For both selection methods in the first training period, the models often underpredicted the higher magnitude blooms.

Many of the forecasts did not predict the exact same CI, or cyanobacterial index, as observed. However in most cases, both methods predicted the same class of bloom as observed. In most cases after increasing the training period to 2013, both ANN models improved their accuracy for predicting the higher magnitude of blooms. The correlation coefficient increased from 0.70 to 0.77 for the loading period selection method and from 0.79 to 0.83 for the Spearman selection method when extending the training period. Both input selection methods had some difficulty in predicting the 2015 HAB because the 2015 bloom was a special case in terms of nutrient loading as well as bloom time. There was a large amount of loading in June and July, which is atypical due to how early it was happening. The 2015 bloom in July was 382 percent larger than any bloom recorded from 2002 to 2014. The monthly discharge for June was the highest recorded and third-highest on record since the United States Geological Survey started collecting data in 1939. Similar to the CART method, the ANN model showed an increase in accuracy when forecasting HABs with an extended training period.

Our method, however does not rely on nutrient variables and instead looks at the growth of algal blooms themselves through measured chlorophyll levels. Our method enables the prediction of severity, location, and distribution of algal blooms, just like both the CART and ANN methods. Because we utilize satellite imagery, however, we are able to simplify the prediction process and increase success. Our algorithm successfully identifies algal bloom components 80 percent of the time, due not only to the neural network but also due to the aforementioned near-certainty with which our satellite images highlight algal blooms.

## Conclusion

Understanding the nature of HABs can lead to their prediction through the data gathered by remote sensing images. We found that not only did the data accurately convey where the HABs would occur, but that the bloom events were predictable. Using a Neural Network to classify the HABs and predict where future HABs would occur, our algorithm was able to render success rates of 80 percent. The future impact of HABs in the Great Lakes can be feasibly lowered by our algorithm, as early response to the HABs can stop their successive propagation through the ecosystem. This can redeem us from our anthropogenic effect on the prevalence of HABs in the Great Lakes, especially Lake Eerie, and can help us keep the Great Lakes a healthy, biodiverse, and oligotrophic ecosystem.

